# Non-linear disease tolerance curves reveal distinct components of host responses to viral infection

**DOI:** 10.1101/113217

**Authors:** Vanika Gupta, Pedro F. Vale

**Affiliations:** Institute of Evolutionary Biology, School of Biological Sciences, University of Edinburgh, EH9 3FL Edinburgh, United Kingdom; Centre for Immunity, Infection and Evolution, University of Edinburgh, EH9 3FL, Edinßburgh, United Kingdom

**Author notes:** Current address: Department of Entomology, 3125 Comstock Hall, Cornell University, Ithaca, NY 14853, USA.

**Keywords:** Infection tolerance, Dose-response curve, Drosophila, Drosophila C Virus, invertebrate immunity, antiviral response

## Abstract

The ability to tolerate infection is a key component of host defence and offers potential novel therapeutic approaches for infectious diseases. To yield successful targets for therapeutic intervention, it is important that the analytical tools employed to measure disease tolerance are able to capture distinct host responses to infection. Here, we show that commonly used methods that estimate tolerance as a linear relationship may be inadequate, and that more flexible, non-linear estimates of this relationship may reveal variation in distinct components of host defence. To illustrate this, we measured the survival of *Drosophila melanogaster* carrying either a functional or non-functional regulator of the JAK-STAT immune pathway (*G9a*) when challenged with a range of concentrations of Drosophila C Virus (DCV). While classical linear model analyses indicated that *G9a* affected tolerance only in females, a more powerful non-linear logistic model showed that *G9a* mediates viral tolerance to different extents in both sexes. This analysis also revealed that *G9a* acts by changing the sensitivity to increasing pathogen burdens, but does not reduce the ultimate severity of infection. These results indicate that fitting non-linear models to host health-pathogen burden relationships may offer better and more detailed estimates of disease tolerance.

## Introduction

Disease tolerance is broadly defined as the host's ability to limit damage and maintain health when faced with increasing pathogen burdens, and is a general feature of host responses to infection [1–5]. Identifying mechanisms that underlie host variation in disease tolerance may therefore offer potentially novel therapeutic targets to treat infections [4, 6–8], an approach already being explored in the context of sepsis [9], HIV [10], influenza [11] and malaria [12]. The key to understanding tolerance is that it cannot be measured by considering host health or pathogen growth separately, but is instead defined by their relationship. This idea is embedded in the original statistical framework of tolerance [13], where it is analysed as a linear reaction norm of host health measured over a range of increasing infectious doses or pathogen burdens. Steep negative slopes for this linear relationship describe groups of hosts that experience a loss in health with increasing loads, while hosts with shallow slopes are able to maintain relatively higher levels of health even as pathogen loads increase, and are therefore relatively more tolerant of infection [1–4].

While the linear reaction norm approach is intuitive and has been useful in advancing the study of infection tolerance (reviewed in [14]), in some cases it may be hindering our ability to achieve a greater mechanistic understanding of the processes underlying host tolerance of infections. For instance, there is no reason to expect the relationship between host health and pathogen burdens to be linear [15], and assuming so may be misleading. For this reason, analytical approaches that allow more flexible, non-linear relationships between host health and pathogen burdens - or ‘tolerance curves’ (Fig 1) - have been proposed [4,7,15]. One advantage of a tolerance curve is that in addition to the rate of health decline with increasing infection loads (the slope), it also allows other health parameters to be estimated, such as host vigour, host sensitivity to increases in pathogen load, and the ultimate severity of infection, which determines how sick a host can get during infection (Fig 1). For example, a recent study of disease tolerance fitted a 4-parameter logistic model to the median survival of *Drosophila* infected with the bacterial pathogen *Listeria monocytogenes*, and this allowed to disentangle changes in fly health during infection that arose due to bacterial pathogenesis and host immunopathology [15], which would not have been possible using classical linear analyses.

**Fig 1.**
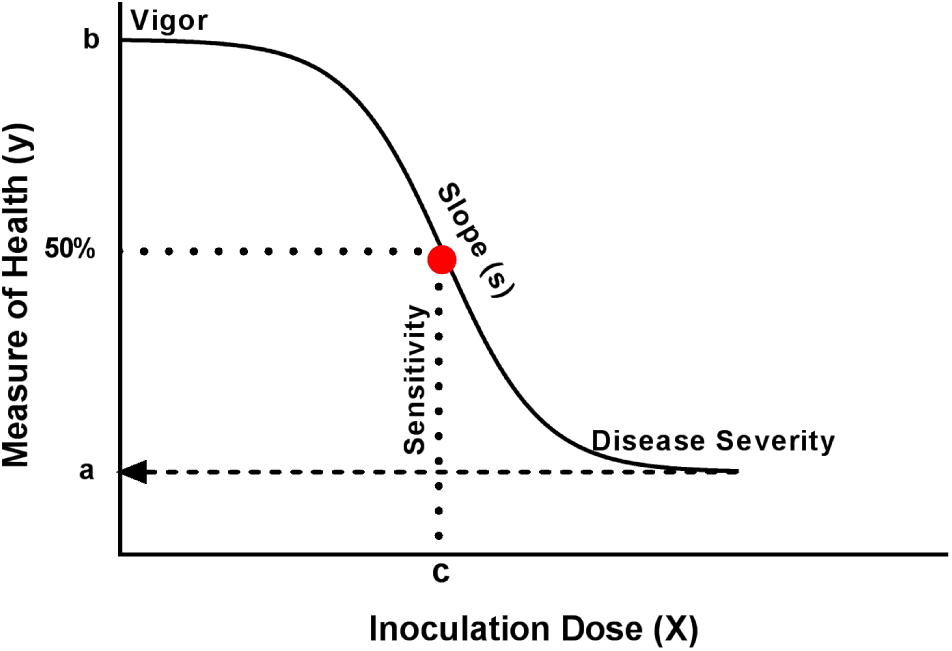
A diagram of how a 4-parameter logistic model can be used to estimate different components of host infection tolerance showing host vigor, the sensitivity, slope, or severity of the dose-response curve [7,15].

Despite these analytical advances, many studies continue to infer tolerance phenotypes from separate measures of host health and pathogen burdens measured at a single infectious dose (for example, [11,12,16]). This approach may provide a general indication that groups of hosts differ in their ability to tolerate infection - for example, when differences in survival are not accompanied by changes in pathogen loads - but they are less useful at describing the rate of health loss with increasing pathogen burdens (the very definition of tolerance), and are also not informative about tolerance at varying infectious doses. This multitude of analytical methods also makes it difficult to draw general conclusions across studies in different species about the ability of hosts to tolerate infection, and how this may vary with genotypic and environmental variation [5,17].

Using both linear and non-linear analyses, here we examine the tolerance response to a systemic viral infection in *Drosophila melanogaster*, which has been used extensively as a model host to dissect the mechanisms underlying tolerance of bacterial and viral infections [15,16,18–20]. *D. melanogaster* infected systemically with Drosophila C Virus (DCV) develops pathology in the reproductive and digestive organs, severe abdominal swelling due to enlargement of the crop and eventually death [21–23]. An epigenetic regulator of the JAK-STAT pathway, *G9a*, was previously identified as a mediator of tolerance to RNA virus infection in *D. melanogaster* [16]. When exposed to a single lethal dose of DCV, fly mutants with a dysfunctional *G9a* showed higher mortality than those with a functional *G9a*, even though there was no difference in the viral loads of the two lines measured at a single time point. This work was notable in providing one of the first examples of immune-mediated tolerance of RNA virus infection, but because it focused on the functional basis of hypersensitivity to DCV, it was designed in a way that did not allow a comprehensive assessment of tolerance. First, infection tolerance was extrapolated from separate analyses of host mortality and viral loads, providing limited information on the role of *G9a* on host vigour, sensitivity to viral growth or the severity of infection that flies can withstand. Further, tolerance was only measured at a single viral dose, making it unclear if the observed tolerance phenotype is dose-specific. Finally, this work only assessed the tolerance phenotype of female flies challenged with a viral infection. Given the prevalence of sexual dimorphism in immunity [24–26], and its expected epidemiological and evolutionary consequences [27,28] it is important to test if infection tolerance may also vary between sexes.

We employed systemic infections in both males and females of two *Drosophila* lines with identical genetic backgrounds, differing only in having a functional or non-functional *G9a* (*G9a*^+/+^ and *G9a* ^−/−^). We challenged these flies with a range of DCV doses and then quantified their tolerance responses using both the slope of linear reaction norm and a non-linear sigmoid model. This allowed us to measure infection tolerance in the most comprehensive way, identifying which components of infection tolerance are affected by a single regulator of fly immunity (*G9a*), while also providing a useful comparison of current methodology to estimate components of infection tolerance.

## Results

### The magnitude of *G9a*-mediated antiviral protection is dose dependent

Following infection with a range of doses of DCV, we found that overall *G9a^−/−^* flies showed significantly higher mortality compared with *G9a*^+/+^(Fig 2a & 2b, Table 1) in line with previously reported effects of this gene on fly survival [16]. However, we found that *G9a^+/+^* and *G9a^−/−^* responded differently to each viral dose, and that the magnitude of the survival benefit of having a functional *G9a* varied with the infectious dose of DCV (Table 1, fly line-by-dose interaction). Notably, mortality was similar between *G9a^+/+^* and *G9a^−/−^* when challenged with the highest dose (10^9^), indicating that the protective effect of a functional *G9a* is no longer observed when flies were challenged with very high doses of DCV. This point highlights the importance of studying infection at a range of doses, as artificially high infectious doses are likely to mask these protective effects.

**Fig 2.**
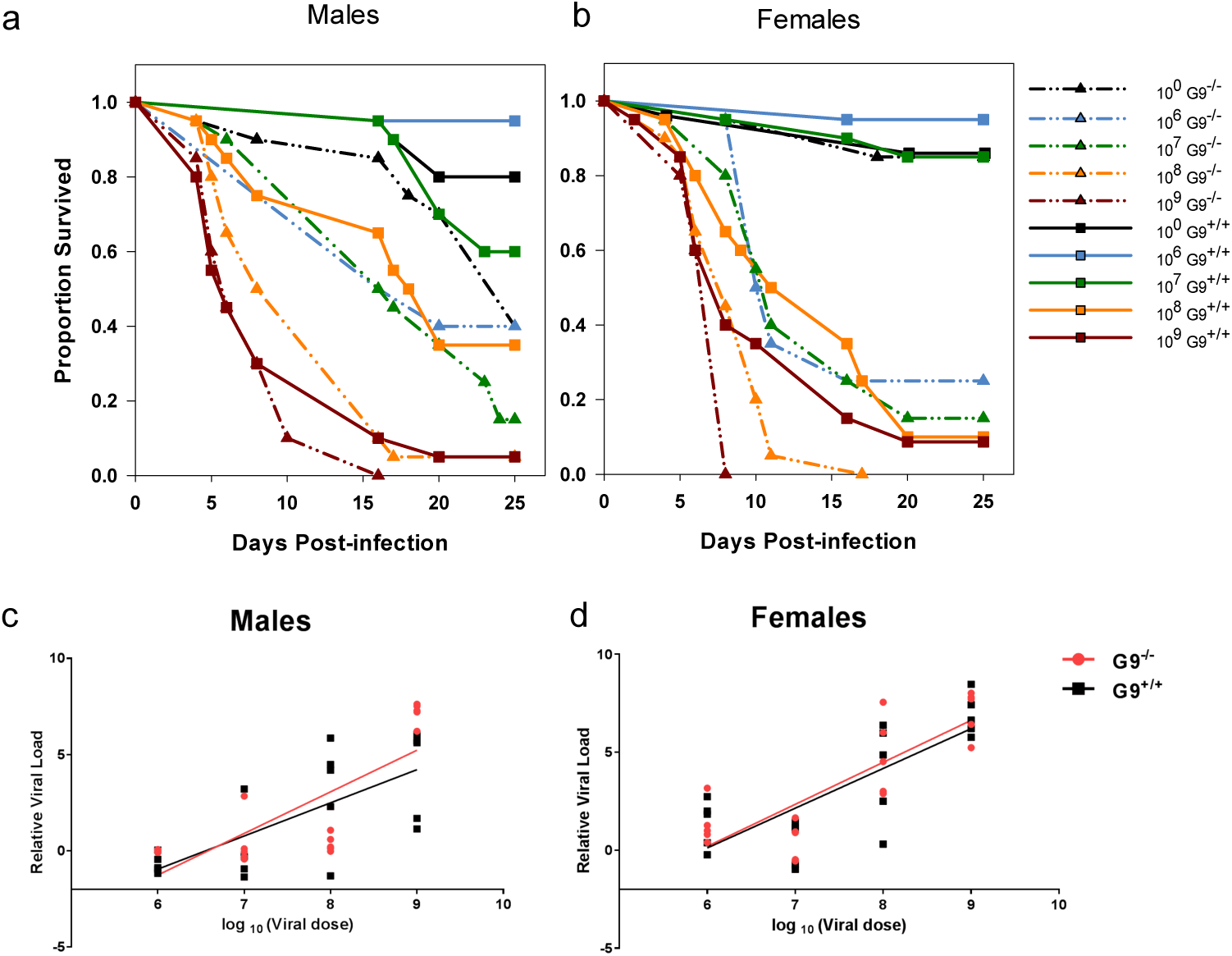
Survival of male (2A) and female (2B) flies challenged with increasing doses of DCV through systemic infection. *G9a^+/+^* flies are shown in solid lines and *G9a^−/−^* mutants are shown in broken lines. Survival was recorded until day 25 post-infection. Each data point is the proportion of 20 individual flies per line / sex / dose combination, and was analysed with a Cox proportional hazard model (Table 1). The DCV titres measured in male (2C) or female (2D) flies exposed to the same DCV doses, 5 days following systemic infection, for *G9a^+/+^* (black) and *G9a^−/−^* (red) lines. Each data point shows the expression of DCV RNA in individual flies relative to the expression of *rp49*, a fly control gene. Lines show there is a significant linear relationship between the dose of DCV flies were challenged with and the viral titre measured after 5 days (details in text and Table 2).

**Table 1:** Cox Proportional Hazards Model

| Source | DF |  | *p* - value |
| --- | --- | --- | --- |
| Fly Line | 1 | 54.81 | <.0001* |
| Dose | 4 | 225.54 | <.0001* |
| Sex | 1 | 0.06 | 0.81 |
| Fly Line × Dose | 4 | 19.79 | 0.0005* |
| Fly Line × Sex | 1 | 0.40 | 0.53 |
| Dose × Sex | 4 | 11.11 | 0.03* |
| Fly Line × Dose × Sex | 4 | 5.33 | 0.26 |

### *G9a^+/+^* and *G9a^−/−^* flies exhibit similar DCV viral loads at all infection doses

Overall, female flies achieved higher viral loads measured 5 days post infection compared to males (Table 2, sex effect). Viral load increased in a dose-dependent manner in both lines (Table 2, Virus dose effect) and did not differ between the two fly lines across all infection doses (Fig 2c & 2d, Table 2, line × dose interaction). These results support previous work using a single dose of DCV, which found that the lower survival of *G9a^−/−^* flies was not attributed to differences in their viral load [16].

**Table 2:**
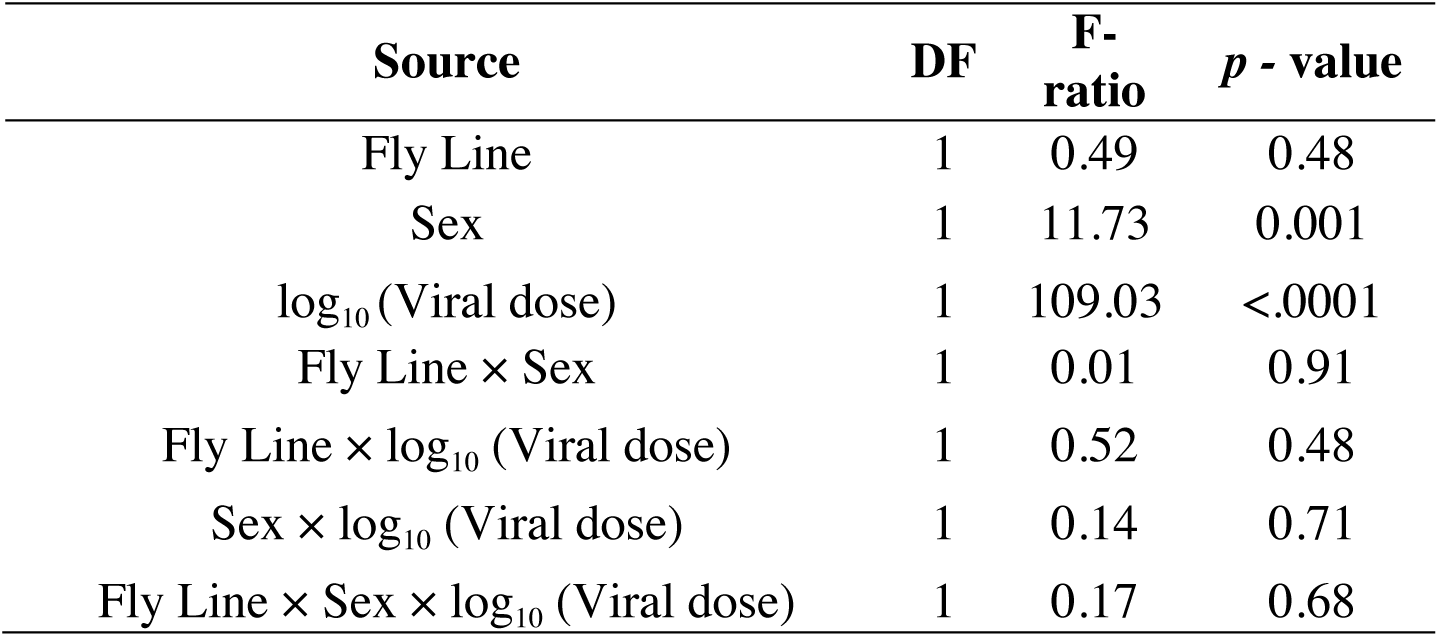
Effect of Fly line, Viral Dose, and Sex on Viral Load

| Source | DF | F-ratio | *p* - value |
| --- | --- | --- | --- |
| Fly Line | 1 | 0.49 | 0.48 |
| Sex | 1 | 11.73 | 0.001 |
| log<sub>10</sub> (Viral dose) | 1 | 109.03 | <.0001 |
| Fly Line × Sex | 1 | 0.01 | 0.91 |
| Fly Line × log<sub>10</sub> (Viral dose) | 1 | 0.52 | 0.48 |
| Sex × log<sub>10</sub> (Viral dose) | 1 | 0.14 | 0.71 |
| Fly Line × Sex × log<sub>10</sub> (Viral dose) | 1 | 0.17 | 0.68 |

### The slope of the linear reaction norm suggests that *G9a*-mediated tolerance is sex-specific

Given that we found a significant positive relationship between DCV infection dose and the viral titre (Males: p < 0.0001, r^2^ = 0.56, Females: p < 0.0001, r^2^ = 0.64; Fig 2c,d), we chose to use viral dose as the measure of pathogen burden in our analysis of tolerance. This approach has the advantage that the covariate (dose) is measured without error, which is one of the assumptions of ANCOVA, and avoids common problems with underestimation of slopes in ANCOVAS when there is experimental variance in the covariate [36]. Differences in tolerance between *G9a^+/+^* and *G9a^−/−^* are indicated by a significant interaction between the viral dose and the fly line for survival, which reflects that the rate at which survival changes with increasing viral doses (tolerance) varies between fly lines. We detected a significant interaction in females (Table 3, Fig 3b), where *G9a^+/+^* females showed higher tolerance (Slope: −1.2 ± 0.4) compared to *G9a^−/−^* females (Slope: −1.8 ± 0.3). In males, however, no significant difference in slopes was detected between *G9a^+/+^* and *G9a^−/−^* (Table 3), even though the survival of *G9a^−/−^* males was lower than thee survival of *G9a^+/+^* males at almost all viral doses (Fig 2a).

**Fig 3.**
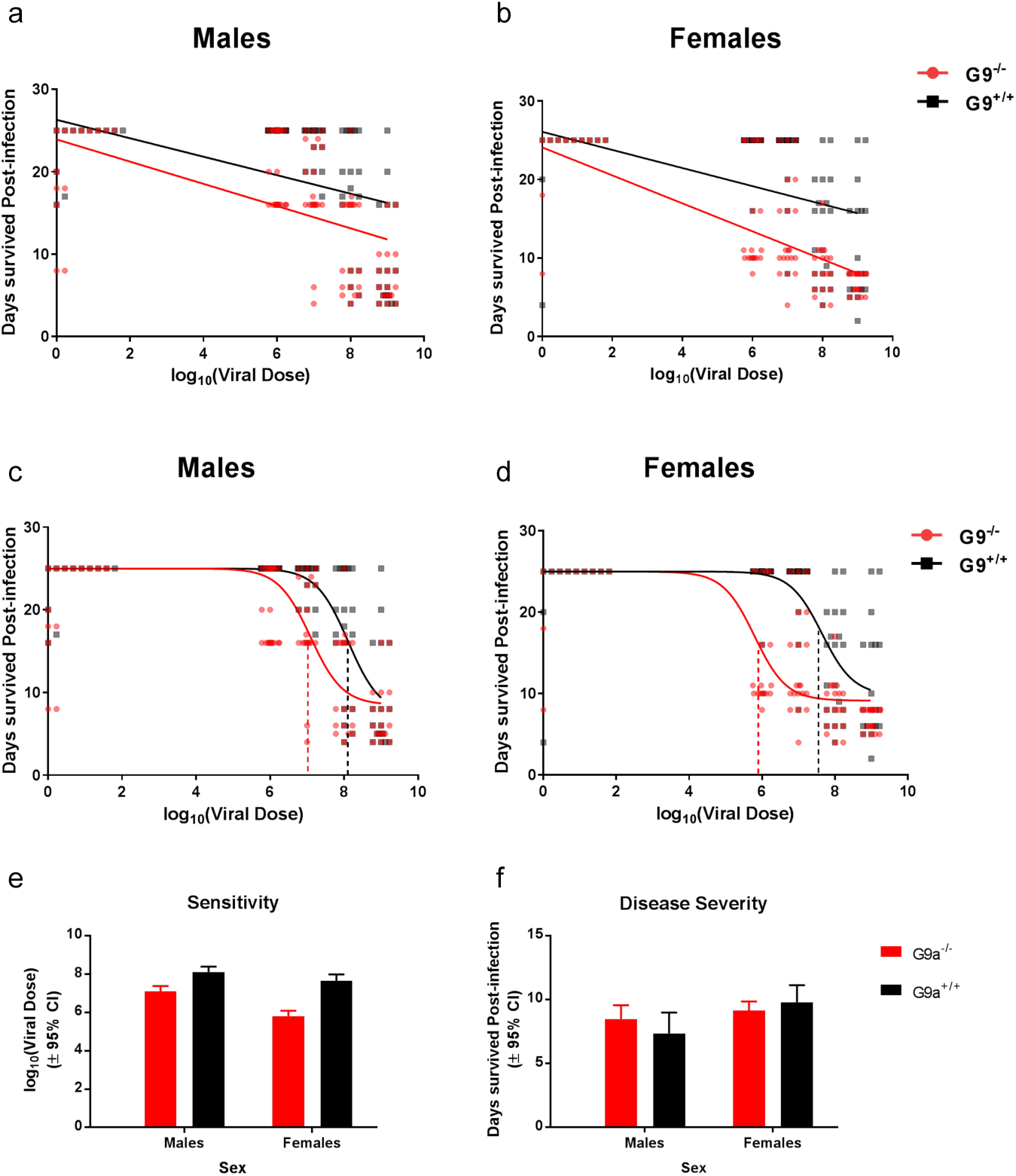
A comparison of linear and non-linear methods to measure infection tolerance in *G9a^+/+^* (black) and *G9a^−/−^* (red) lines. The relationship between host survival and viral dose analysed using linear models for males (2A) and females (2B) or non-linear 4-parameter logistic models for males (2C) and females (2D). Each plot shows individual data for 20 individual flies per line / sex/ dose combination. Fig 2E and 2F show the mean and 95% confidence intervals for sensitivity and ultimate severity, respectively, extracted from the non-linear models. Significant pairwise differences are indicated with an asterisk and linear and non-linear model details are reported in the main text and in in Tables 3 and 4.

**Table 3:**
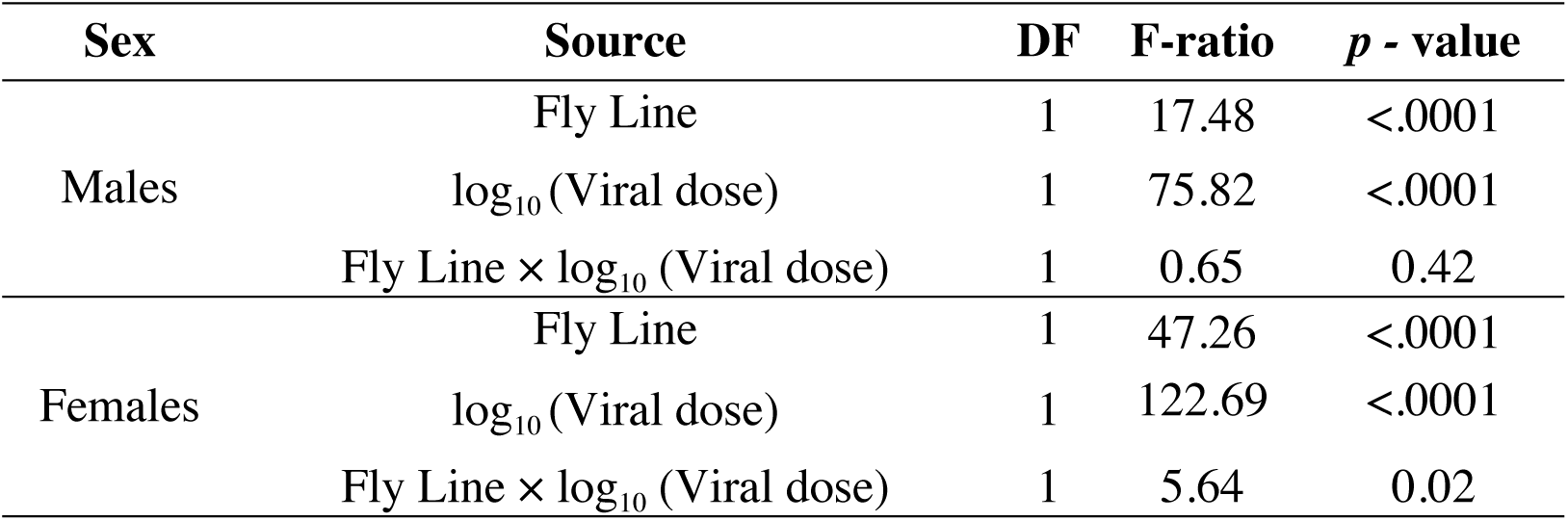
Linear tolerance: General Linear Model studying effect of Fly line and Viral Dose on post-infection survivorship in males and females

### Non-linear models reveal that *G9a* affects the sensitivity to infection but not its severity

We then analysed the relationship between fly survival and viral dose using a non-linear 4-parameter logistic model [7,15,33]. Using this non-linear model allowed us to assess how *G9a* affected the sensitivity of flies during infection and the ultimate severity of DCV infection in both functional and deficient versions of this regulator of the JAK-STAT pathway (Fig 1). A comparison of the overall fit of the curves showed that *G9a^+/+^* and *G9a^−/−^* backgrounds have distinct tolerance profiles during DCV infection (p < 0.0001, Males: F_2, 196_ = 14.8, Females: F_2, 196_ = 50.71). In contrast to the analysis assuming a linear relationship, we found that both male and female *G9a^−/−^* flies differed from *G9a^+/+^* in their ability to tolerate DCV (Fig 3c & 3d, Table 4). This non-linear analysis showed that *G9a^+/+^* flies have a significantly higher inflection point compared to *G9a^+/+^* flies, suggesting that *G9a*^+/+^ flies are less sensitive (or more tolerant) to increasing viral doses. The sensitivity to infection differed by 70-fold between *G9a^+/+^* and *G9a^−/−^* in females (p<0.0001) while we detected a ten-fold difference in males (p<0.0001) (Fig 3e). We did not find any difference between *G9a^+/+^* and *G9a^−/−^* in the severity of infection (Table 4, Fig 3f). Our results therefore show that *G9a* regulates the sensitivity to infection, accelerating the onset of infection-associated mortality when it is dysfunctional, without causing substantial effects on the disease severity ultimately experienced by infected flies.

**Table 4.**
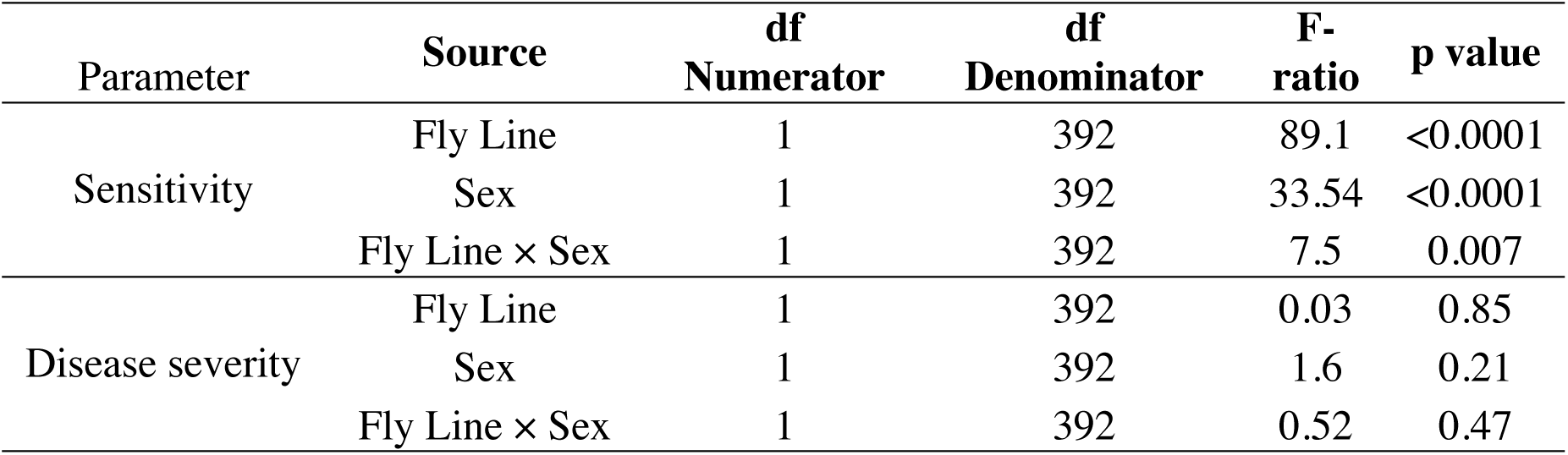
Parameters of a 4-parameter logistic model: Effect of Line, Sex, and Line-by-Sex interaction

### Lack of *G9a* leads to sex-specific expression of the JAK-STAT ligand *upd3*

The differences between *G9a^−/−^* males and females in the sensitivity to increasing viral loads (Fig 3E) prompted us to investigate if this may be due to sex-specific regulation of fly immunity by *G9a.* As *G9a* is know to be an epigenetic regulator of the JAK-STAT pathway, we measured the expression of JAK-STAT pathway genes in flies receiving the highest viral concentration, 5 days following DCV infection (the same day on which viral loads were quantified). Compared to flies with a functional *G9a, G9a^−/−^* mutants showed a significant increase in the expression of the JAK-STAT ligand *upd3*, and this effect was stronger in male flies (Fig 4; Table 5). Males showed generally higher expression of the JAK-STAT receptor *domeless*, although this effect was independent of *G9a* status (Fig 4; Table 5). *G9a^−/−^* flies also showed significantly higher expression of the negative regulator of JAK-STAT, *socs36E*, but this effect did not differ between males and females. Finally, we found no effect of either sex or *G9a* on the expression or *turandotA* (*totA*), which is commonly expressed in response to stress.

**Fig 4.**
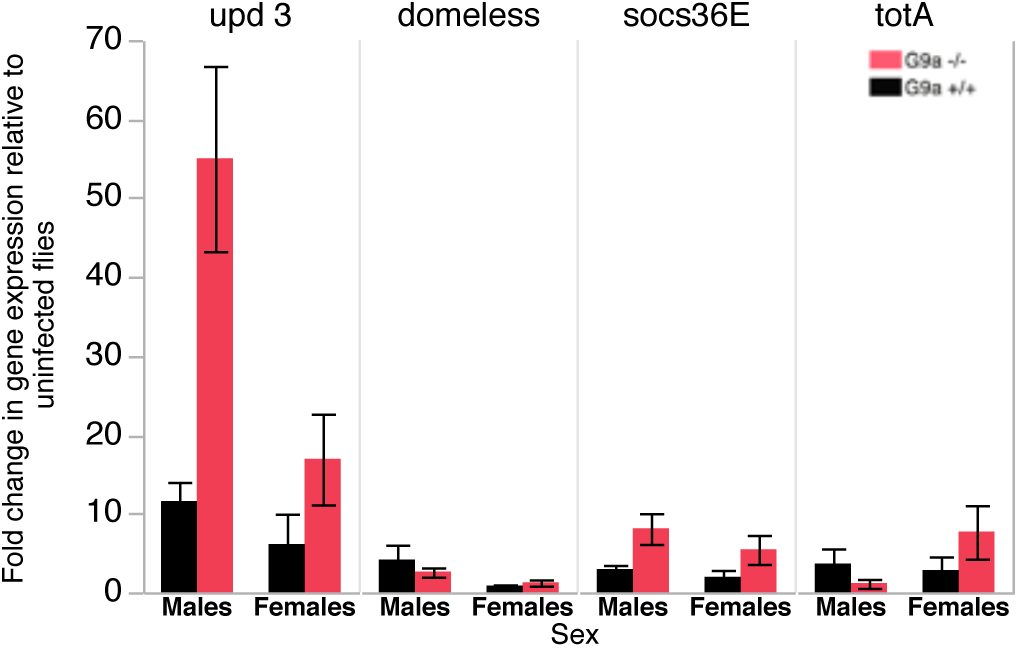
Expression of JAK-STAT immune genes. Gene expression relative to the internal control gene *rp49* was quantified in five replicate individual male and female flies flies exposed to the highest dose treatment (10^9^) relative to their expression in uninfected controls. Data show mean±SE. See table 5 for statistical analysis output.

**Table 5.** Effect of fly line and sex on JAK-STAT immune gene expression

| Gene | Source | DF | F Ratio | p-value |
| --- | --- | --- | --- | --- |
| *upd 3* | Sex | 1 | 8.362 | 0.012 |
|  | Fly Line | 1 | 12.995 | 0.003 |
|  | Fly Line × Sex | 1 | 4.712 | 0.048 |
| *domeless* | Sex | 1 | 5.101 | 0.038 |
|  | Fly Line | 1 | 0.295 | 0.595 |
|  | Fly Line × Sex | 1 | 0.897 | 0.358 |
| *socs36E* | Sex | 1 | 1.577 | 0.227 |
|  | Fly Line | 1 | 9.009 | 0.009 |
|  | Fly Line × Sex | 1 | 0.343 | 0.566 |
| *totA* | Sex | 1 | 1.738 | 0.206 |
|  | Fly Line | 1 | 0.306 | 0.588 |
|  | Fly Line × Sex | 1 | 2.883 | 0.109 |

## Discussion

Targeting mechanisms that promote greater tolerance of infection is a promising addition to our current arsenal of strategies to fight infection [3, 6–8]. However, if this is to be a successful undertaking, it is crucial that the analytical tools we use to measure tolerance are able to capture distinct host responses during infection. A major aim of this study was to evaluate two analytical methods to measure tolerance (linear reaction norms and non-linear curves), in order to assess the benefit and limitations of each approach.

Differences in infection tolerance between groups of hosts are commonly extrapolated from single infection doses and by assessing host survival and pathogen burdens separately [11,12,16]. These experiments are useful in detecting the effect of specific mechanisms underlying host tolerance, but the limitations of single-dose tolerance experiments arise because a host's ability to limit the damage, and therefore tolerate infection, is not necessarily independent of the within-host pathogen burden it suffers. The approach arising from the evolutionary ecology of infection, which measures tolerance as a linear reaction norm, provides additional information, such as an estimate of general vigor from the intercept, while its slope gives an estimate of the rate of decline in the health [1,14]. While useful, there are at least two limitations to interpreting infection tolerance as a linear reaction norm. First, there is no biological requisite for the relationship between pathology and pathogen burdens to be linear [10,37]. Second, there are statistical caveats associated with measuring differences between linear slopes when data ranges do not overlap, which is likely if hosts also differ in the mean and variance of their pathogen loads (the x-axis of the reaction norm) [38,39].

In this regard, using non-linear sigmoid models offers greater flexibility to fit a range of possible relationships (including a linear approximation) between host health and pathogen loads [15]. Moreover, non-linear tolerance curves provide additional information on several components of host responses to infection, such as the sensitivity and severity of infection. Differences between groups of hosts in any of the parameters extracted from a non-linear model (slope, sensitivity, severity) may therefore reflect distinct underlying mechanisms that either promote greater damage prevention or increase damage repair during infection [15]. Employing such methodology to a range of host-pathogen systems may therefore yield useful targets for therapeutic interventions that increase host disease tolerance [7].

Despite all the stated advantages, one potential drawback of using non-linear sigmoid models to study tolerance is that the slope of health decline is measured over a very small range of pathogen doses. Accurate estimates of that slope would therefore require many estimates of host health within a very narrow range of pathogen burdens, which may be experimentally challenging. For this reason, in the current analysis we fixed the slope to −1, which also increased the statistical power to estimate the two parameters of interest (sensitivity and severity). In this respect, we propose that using the linear reaction norm approach could be useful if the main variable of interest is the rate at which hosts lose health, which is estimated along the full range of pathogen burdens. Non-linear approaches are more informative if the parameter of interest is the dose that causes the greatest shift in host health, or if the question relates to how hosts may differ in the ultimate severity of an infection.

A subsidiary interest of this work was to quantify *G9a*-mediated tolerance of DCV in both male and female *Drosophila.* Both linear and non-linear models consistently showed that *G9a^+/+^* females have higher tolerance than *G9a^−/−^* females, when measured across a range of DCV doses. However, the two models presented different results in the case of males. Applying the classical linear reaction norm approach, the linear slope showed no difference in the rate of survival with increasing DCV doses, while the non-linear fit indicated significant differences between *G9a^+/+^* and *G9a^−/−^* males in their sensitivity to infection. Further analysis of the sensitivity and severity of infection parameters extracted from the non-linear model also showed that the effect of *G9a* in mediating host sensitivity to increases in DCV dose was greater in females. It is important to note that while previous work also found *G9a^−/−^* females were hypersensitive to DCV infection, this was only assessed by measuring survival at one dose and infection tolerance was not measured in male flies [16].

While males and females are generally susceptible to the same pathogens, sexual dimorphism in immunity is present in a wide range of species [24,40,41], and sex differences in infection tolerance are documented for all classes of viral, bacterial, fungal and parasitic infections [see [27] for a review]. Differences between males and females in the ability to tolerate infection will directly impact on the pathogen loads within hosts, and as a consequence, also affect host shedding of pathogen transmission stages [42,43]. Sexually dimorphic tolerance is therefore predicted to generate potentially important heterogeneity in pathogen spread and evolution [27].

The mechanisms underlying sex-specific *G9a*-mediated effects on infection tolerance are not clear. *G9a* is a histone methyltransferase [44,45], and the protective effect of *G9a* during viral infection has been previously shown to be driven by the regulation of the JAK-STAT pathway [16]. Specifically, *G9a* is known to alter the methylation state of the positive regulator *domeless*, the negative regulator *Socs36E* and downstream pathway components (*TotA, vir-1*) of the JAK-STAT pathway [45]. Fly mutants without a functional *G9a* show increased mortality due to immunopathology caused by excessive expression of these downstream genes [16]. Notably, *G9a, domeless* and its ligands *upd1, upd2* and *upd3* are all found on the X-chromosome in *Drosophila* [46,47]. We hypothesised that these X-linked regulators of fly innate immunity could underlie the sexually dimorphic tolerance response we observed. However, while we detected sex differences in the expression of *domeless* (Fig 4), these effects were not a consequence of *G9a* function. Instead we found the largest sex-specific effect on the expression of the *upd3* (Fig 4). Previous work had shown that upd3 expression is higher *G9a*^−/−^ flies, resulting in immunopathology [16]. Our work shows that these effects differ between sexes and are especially strong in males.

In summary, we show that *G9a* mediates tolerance during infection with DCV over a large range of viral doses and that this response differs between males and females. Our study therefore places emphasis on the importance of incorporating both males and females in studies of immunity. Our results also stress that conclusions about disease tolerance will vary according to the method used to estimate the relationship between host health and pathogen burdens. We suggest that a combination of linear and non-linear models is ideal to achieve estimates of the rate of decline in health following infection but also of subtler components of hosts’ responses, such as the sensitivity to increases in pathogen loads and the ultimate severity of disease experienced during infection. Further, these methods are applicable not only to the declines in health experienced during infection, but could in principle be applied to other diseases, such as cancer [48]. In general, our understanding of host responses to disease requires a more complete assessment of resistance and tolerance mechanisms across a range of genetic and environmental contexts.

## Methods

### Fly stocks

Experiments were conducted on *Drosophila melanogaster* mutants *G9a^+/+^* and *G9a^−/−^* (also known as *G9a^DD2^*), kindly provided by R. van Rij (Radboud University, Nijmegen, NL). *G9a^−/−^* was originally constructed by excision of P-element *KG01242* from the 5’ UTR of the gene [16]. As a control for the mutant line, a P-element excision line was generated in the same genetic background that restored the functional phenotype of *G9a*, referred to as *G9a*^+/+^. Both fly lines were maintained on standard Lewis Cornmeal medium under standard laboratory conditions at 25°C, 12h: 12h Light: Dark cycle.

### Generation of experimental flies

To set up the experiment, we collected eggs from 15 males and 15 females of each line, kept in vials containing 6mL Lewis medium supplemented with dry yeast to encourage egg laying. We set up ten replicate vials per genotype. Flies were left to oviposit in the vials for 24 hours before being removed and allowing eggs to develop under standard rearing conditions. To control the larval density of these flies, egg density was maintained between 80-100 eggs per vial by removing excess eggs when required. Flies emerging from these vials were challenged with DCV and their mortality post – infection and viral load were measured.

### Virus preparation and titration

*Drosophila* C virus (DCV) is a ssRNA virus of the family *Dicistroviridae.* We obtained a viral stock by amplifying DCV in Drosophila S2 cells as described previously [29]. Cell homogenate containing DCV was passed through a sucrose cushion, and the resulting pellet was suspended in 10mM Tris-HCl (pH 7.3). The virus stock was stored in small aliquots at - 80°C until further use. To estimate the viral infection dose, we measured the absolute quantity of virus in the stock culture using quantitative Real-Time PCR (qRT-PCR). Briefly, an aliquot of this virus stock was taken to extract RNA using TRI reagent. This RNA was further used in RT-PCR to obtain cDNA, and this cDNA was then serially diluted 10-fold until dilution 10^-10^. SYBR Green-based quantitative Real-time PCR (qPCR) on these samples was performed, and the first dilution where no viral cDNA was detected was taken as zero, and stock viral quantity was back calculated from this reference point. Using this method, viral copy number in the stock was estimated to be around 10^9^ viral copies per *μ*L.

### Virus infection

We exposed experimental flies to 5 viral doses – 0 (control), 10^6^, 10^7^, 10^8^, and 10^9^ DCV copies, obtained by diluting the viral stock with 10mM Tris-HCl (pH 7.3). Flies were infected systemically by intra-thoracic pricking with a needle immersed in DCV suspension under light CO_2_ anesthesia. The effect of injury caused by pricking was controlled by including sham-infections performed using a needle dipped in sterile10mM Tris-HCl (pH 7.3).

### Experimental set-up

#### (a) Post-infection survival

3-4-day-old adults were systemically infected by intra-thoracic pricking with DCV. For each Fly line × Sex × Dose combination, we infected 20 replicate individual flies (400 flies in total). Following infection, flies were housed individually its own vial and flies were monitored daily for mortality. Flies were transferred onto fresh medium once a week. Survivorship was followed for 25 days post-infection and flies that were still alive at the end were censored in the analysis.

#### (b) Viral Load

An additional five individuals for each Genotype × Sex × Dose combination were infected as described above to quantify differences in viral growth following infection, using the expression of DCV RNA. Flies were individually placed in TRI reagent (Ambion) following five days of infection (5 DPI) and stored at −80°C. Total RNA was extracted from these flies homogenized in Tri-Reagent (Ambion), which includes a DNase treatment step, reverse-transcribed with M-MLV reverse transcriptase (Promega) and random hexamer primers, and then diluted 1:2 with nuclease-free water. qRT-PCR was performed on an Applied Biosystems StepOnePlus system using Fast SYBR Green Master Mix (Applied Biosystems) and DCV primers DCV_Forward: 5’ AATAAATCATAAGCCACTGTGATTGATACAACAGAC3’,DCV_Reverse: AATAAATCATAAGAAGCACGATACTTCTTCCAAACC). The expression of DCV transcripts was normalized to transcript levels of the housekeeping gene *rp49* (Dmel_rp49 Forward: 5'ATGCTAAGCTGTCGCACAAATG 3’; Dmel_rp49 Reverse: 5’ GTTCGATCCGTAACCGATGT 3’). and expressed as fold change relative to the control flies, calculated as 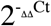 [30].

#### (c) Gene Expression

Using the same cDNA samples described above, we measured the expression of immune genes likely to be affected a *G9a* deletion, to test for genotype-specific and/or sex-specific differences upon infection. Given that G9a is a regulator of the JAK-STAT pathway, we focussed on measuring the expression of the extracellular ligand *upd3*, it's transmembrane receptor *domeless*, and downstream regulators *socs36E*, and *totA*. Gene expression was quantified in flies exposed to the highest dose treatment (10^9^) relative to their expression in uninfected controls, following the procedure decribed above for viral loads.

### Statistical Analysis

To analyze differences in survival of each host line according to the virus dose they were challenged with, we analyzed post-infection survival data using a Cox proportional hazards model with ‘Fly line’, ‘Dose’ and ‘Sex’ and their interactions, fitted as fixed effects. To assess differences in viral titers following infection with infection doses, we analysed the Log_10_ viral titer measured at 5DPI using a linear and fitted ‘Fly line’, ‘Dose’ and ‘Sex’ and their interaction as fixed effects.

We tested for differences in infection tolerance between fly lines in two ways. First, we use the classical approach and analyzed tolerance as a linear reaction norm [19,31,32]. A general linear model was fitted separately for males and females to study the effect of ‘viral dose’ and ‘fly line’ on fly health (DPI). In this analysis ‘viral dose’ was treated as a continuous covariate and ‘fly line’ as a categorical factor. Given that infection tolerance refers to the rate at which hosts lose health with increasing viral loads, the analysis of interest is how the slope of the two fly lines differs, that is, if survival is significantly affected by the ‘fly line-by-viral load’ interaction.

As a second approach to quantify infection tolerance, we fitted a non-linear 4-parameter logistic model (Eqn. 1) which is commonly used to assess dose-response curves [15,33,34] to the survival and virus dose data. Using GraphPad Prism v6.0, we fitted the following equation:

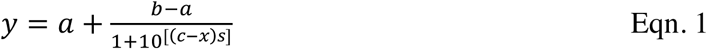

where *a* is the level of disease severity, *b* is a host's general vigour, *c* is the sensitivity to increases in pathogen dose (in this case, the pathogen dose that results in the level of health halfway between the min and max health, similar to EC_50_), *s* is the slope of the logistic model, and *x* is the pathogen dose. Since the experiment was terminated 25 days post-infection, we constrained the upper limit of the model (host vigour) to 25 (DPI) (see also [15]). To test if these tolerance curves differed between fly lines, we compared the overall fit of the curves using extra sum-of-squares F-test [35]. Using these fitted models, we also examined if lines differed in their sensitivity and disease severity, after fixing the slope to −1. To test for differences in the severity and sensitivity to infection between male/female groups, or between *G9a* lines, we carried out two-way ANOVA with Line and Sex as fixed effects, using Tukey's HSD as a post-hoc analysis of pairwise comparisons. Unless otherwise stated, all analyses were carried out in JMP 12 (SAS).

## Acknowledgements

We thank J. Oliveira and T. Little for comments on the manuscript. We thank D. Obbard and lab members (Edinburgh) for general technical support and for advice about DCV infection; we are also grateful to R. van Rij (Nijmegen) for providing the fly lines used in this study. Finally, we thank H. Cowan and H. Borthwick for help with media preparation, and K. Monteith general technical support.

